# Single-nucleus transcriptome analysis provides new insights into B chromosome elimination in sorghum

**DOI:** 10.1101/2025.10.07.680864

**Authors:** T Bojdová, P Abaffy, K Holušová, L Valihrach, M Karafiátová, J Bartoš

## Abstract

B chromosomes (Bs) are supernumerary entities found in many plant species, with some exhibiting tissue-specific elimination. In *Sorghum purpureosericeum*, extensive level of the B chromosome elimination occurs during embryogenesis. It progresses quickly and affects most of the embryonic organs, leaving the B chromosome maintained mainly in limited regions of meristems. Dynamic of the process and rarity of the transcripts associated with elimination make its capturing challenging. To address this, we performed single-nucleus RNA sequencing (snRNA-seq) on embryos actively undergoing B chromosome elimination. The snRNA-seq approach enabled the detection of a greater number of B-linked transcripts compared to previous method. Among all, we identified the nuclei with B-specific transcripts, which prevalently clustered together forming cluster 10 and contributing to cluster G2/M. The detailed analysis of cluster 10 revealed two distinct subpopulations of B-containing nuclei with divergent transcriptional profiles. The genes expressed in one subpopulation (sCL10-1) indicated that elimination of the B chromosome might be a factor for the split of the two subpopulations; namely that nuclei in sCL10-1 prepare for / undergo elimination, while the other subpopulation is expected to exhibit regular segregation of the B chromosome. Our analysis shows a direction so far missing in current studies and highlights a clear benefit of the single-cell approaches for studying specific behaviour of the B chromosomes.

## Introduction

B chromosomes (Bs) are dispensable supernumerary elements found in about 3000 species, generally existing alongside the standard A chromosome set in all tissues (D’Ambrosio et al., 2017). However, in some species, Bs are eliminated in a tissue-specific manner (Nygren, 1957; Baenziger, 1962; Mendelson & Zohary, 1972; Mochizuki, 1957; Karafiátová et al., 2024). Their elimination patterns are highly variable; e.g *Poa alpina* and *Agropyron cristatum* lose Bs from adventitious roots but retain them in primary roots (Baenziger, 1962) whereas *Aegilops speltoides*, *Aegilops mutica*, and *Poa timoleontis* eliminate Bs from primary roots (Nygren, 1957; Mochizuki, 1957; Mendelson & Zohary, 1972). A particularly striking case is *Sorghum purpureosericeum*, where Bs are nearly completely lost from mature somatic tissues (Karafiátová et al., 2024), similar to germline-restricted chromosomes (GRCs) in songbirds, which are absent from all somatic cells (Hodson et al., 2021; Borodin et al., 2022).

The onset of B chromosome elimination can be traced back to embryogenesis, as seen in *Aegilops speltoides* (Ruban et al., 2020) and *Sorghum purpureosericeum* (Karafiátová et al., 2024)(Bojdová et al., 2025). Bs are progressively eliminating during embryonic development and therefore these chromosomes are inherently programmed to persist only in specific tissues. Recent studies in *A. speltoides* (Boudichevskaia et al., 2020) and *S. purpureosericeum* (Bojdová et al., 2025) focused on identification of candidate genes linked to B chromosome elimination in embryo transcriptome. However, these studies have relied on whole embryo RNA-seq, which conceals cell-type-specific dynamics critical to understanding the elimination process. As embryo comprises a heterogeneous mixture of cells and only a small subset undergoes B chromosome elimination, transcripts associated with it are likely to be underrepresented in conventional transcriptomic experiments. In whole embryo RNA-seq, the candidate transcripts tend to fade in the transcriptomes of vast majority of cells (either with stable B chromosome or after its elimination) and it is impossible to assign them to the specific cells undergoing elimination. Single-cell or single-nucleus RNA sequencing (sc/snRNA-seq) could offer a solution to this limitation by allowing the detection of rare cell populations undergoing elimination and corresponding transcripts. To date, no single-cell transcriptomic study targeting B chromosome elimination has been conducted, largely due to the plant species pioneering the B chromosome research have Bs stably maintained across cell types. Further, this approach has neither been used for germline-restricted chromosomes (GRCs) eliminated from somatic tissues, presenting a compelling parallel to B chromosome elimination. GRCs are much more frequent than B chromosomes undergoing elimination (Hodson 2021, Borodin et al., 2022), but challenging to study due to the difficulty of obtaining germline tissues at the appropriate developmental stages. Thus, *Sorghum purpureosericeum* offers a possibility to be model system to investigate the molecular mechanisms underlying programmed chromosome elimination.

In this proof-of-concept study, we explore whether single-nucleus RNA sequencing is beneficial to uncover cell-to-cell transcriptional heterogeneity associated with B chromosome elimination. By profiling embryonic tissue of *S. purpureosericeum* at 21 days after pollination (DAP), we aimed to determine whether elimination-specific signatures can be resolved in the population of cells possessing B chromosomes. Although exploratory, our work highlights the potential of this approach to uncover cell-specific mechanisms governing chromosome fate. The findings presented lay the groundwork for future studies and establish a conceptual framework for using sc/snRNA-seq to investigate B chromosome–specific phenomena at the single-cell resolution.

## Results and Discussion

### Single-nucleus RNA-seq offers a promising strategy for solid tissues resilient to protoplasting

To address the changes in the cells of wild sorghum *S. purpureosericeum* embryos, which direct the B chromosome to its elimination, we investigated the RNA-landscape at the resolution of single nuclei. We analysed the nuclei of the embryos of approx. 21 DAP (Fig. 1A, B), where the extent of the B chromosome elimination accompanying the embryo differentiation was documented (Fig. 1C, Karafiátová et al., 2024; Bojdová et al., 2025). At targeted stage, the B chromosome elimination significantly proceeded, and many embryonic tissues are free of B chromosome. However, even cells showing signal from B-specific probe do not exhibit stable presence of the B chromosome and its elimination continued as evidenced by the presence of micronuclei (Fig. 1C, D). Embryonic nuclei suspension is thus a complex cocktail containing the nuclei lacking B chromosome, harbouring the B chromosome and undergoing the B chromosome elimination (Fig. 1E).

**Figure 1:**
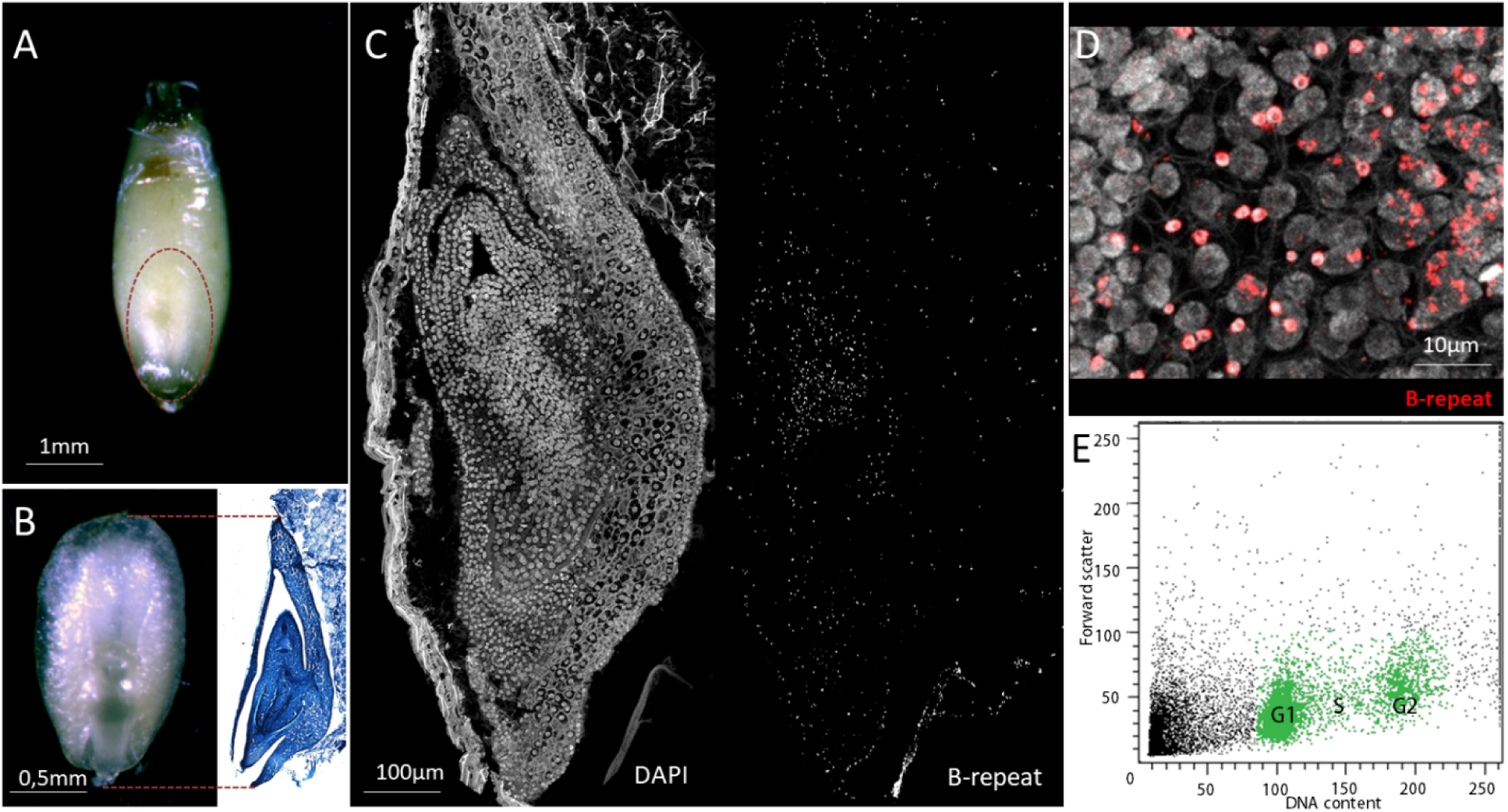
Embryo anatomy and progress of the B chromosome elimination in wild sorghum *S. purpureosericeum*. **A**. Developing seed of wild sorghum approx. 21DAP. Embryo position is indicated by red-dashed line. **B**. Intact isolated embryo of 21 DAP (ventral side) and its transversal section stained with toluidine blue illuminating embryo organogenesis. **C.** Distribution of the B chromosome in developing embryo visualized using FISH with B-specific probe cocktail on embryo cryo-sections; DAPI counterstaining (left) and signal from B-repeat cocktail (right). **D.** Chimeric composition of embryonic tissue consisting of three types of nuclei – harbouring B chromosomes (positive for B-repeat signal), undergoing B chromosome elimination (B chromosome located in micronucleus) and yet free from B chromosome (nuclei absent for specific signal). B chromosome is detected using FISH with B-repeat cocktail (red). **E.** Nuclei isolation for snRNA-seq using flow cytometry. Green-marked populations represent the mix of +B and 0B embryonic nuclei in G1/S/G2 phase used as input DNA for sequencing library.

A total of 100,000 nuclei were collected through flow sorting (Fig. 1F). The nuclei formed two distinct populations in flow cytometric analysis differing in ploidy level corresponding to cell cycle phases (G1 and G2) and not by the B chromosome presence/absence (Fig. 1F). Both population as well as nuclei with intermediate DNA content (S phase) were profiled via droplet-based snRNA-seq. After data filtering, 5,214 nuclei (representing approx. 30% recovery) were retained for further analysis. On average, 1,379 genes per nucleus were detected, with a mean read depth of 19,655 reads mapped to genes and an average unique molecular identifier (UMI) count of 2,137 per cell. In total, expression of 35,745 genes was recorded within all cells, representing an improvement over the 30,580 genes reported in the previous RNA-seq study (Bojdová et al., 2025). Importantly, detection of expressed B chromosome genes also increased, from 2,012 to 3,024.

Candidate genes controlling the B chromosome elimination in *S. purpureosericeum* have been identified previously (Bojdová et al., 2025). However, the analysis was conducted on whole embryos, disabling the spatial tracking candidate gene expression. In contrast, single-cell RNA sequencing (scRNA-seq) would enable such resolution, nevertheless protoplasting embryo samples for scRNA-seq presents technical challenges. To circumvent these difficulties, we opted for single-nucleus RNA sequencing (snRNA-seq), which sequences nuclear transcripts, thereby eliminating the need for protoplasting. This approach has been successfully applied in plant embryo transcriptomics studies (Kao et al., 2021; Long et al., 2021). In both studies, the reported gene detection metrics were comparable with those obtained in our experiment. Considering the pros and cons of the method, the RNA-analysis at the single nuclei resolution represents a meaningful compromise yet providing valuable knowledge. Our trial experiment shows a clear benefit of this method and outlines a new trend in the study of B chromosome elimination.

### Functional assignment of snRNA-seq clusters identified population of nuclei harbouring the B chromosome

Unsupervised clustering analysis using Seurat software (Satija et al., 2015) identified distinct embryonic cell populations based on their nuclear transcriptomic profiles. Visualization of the data via uniform manifold approximation and projection (UMAP) (Becht et al., 2019) revealed thirteen cell populations at the 0.5 resolution (Figure 2A), with cluster sizes ranging from 96 cells (cluster 12) to 906 cells (cluster 0). A total of 1,710 genes were assigned as marker genes (using default settings) distinguishing these 13 snRNA-seq clusters (Figure 2B, Source Data 1). To annotate clusters, we firstly took advantage of published embryonic single-cell transcriptomes of barley (Peirats-Llobet et al., 2023) and maize (Wu et al., 2025) comparing *S. purpureosericeum* marker genes to their homologs in these datasets. Additionally, we searched for marker genes in scPlantDB (He et al., 2024), a database of scRNA-seq studies, and examined the previously characterized biological functions of the identified cluster-specific marker genes (Source data 2). This strategy enabled the classification of four major nuclei groups comprising 10 out of 13 clusters: embryo (emb-1 to emb-5), scutellum (scu), endosperm (end-1, end-2) and actively proliferating cells (S, G2/M).

**Figure 2:**
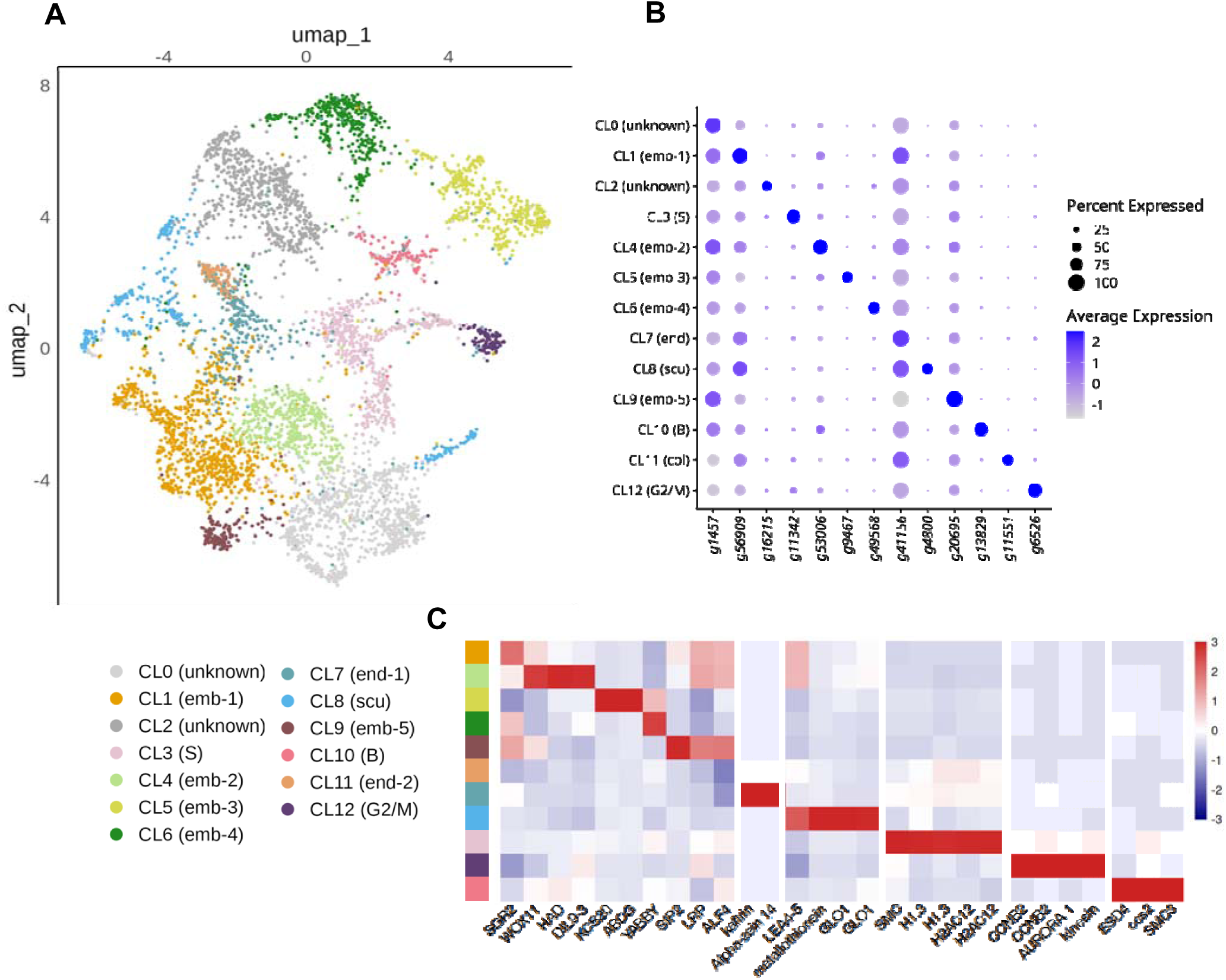
Single-nucleus RNA-seq clustering and cluster annotation of *Sorghum purpureosericeum* embryo tissues. **A.** UMAP visualization of 13 snRNA-seq clusters from 5,214 cells. Each dot indicates a single nucleus. **B.** Expression dotplot of cluster-specific top marker genes for each cell type. Dots size highlights proportion of cells expresing given marker genes in each cluster and dots color corresponds to mean expression in all cells in given cluster. **C** Heat map of average SCT-transformed expression of annotated marker genes across clusters. First column represent color codes for the clusters identical to panel A.

Among clusters assigned to the embryo, CL1 (emb-1) expressed *SHOOT GRAVITROPISM 2 and 6 (SGR6*), genes associated with root gravitropism (Kato et al., 2002) as well as *WUSCHEL-related homeobox 11 (WOX11)*, which is linked to root apical meristem (RAM) function (Zhao et al., 2009). CL4 (emb-2) included markers of vascular tissue, such as a phloem-associated *haloacid dehalogenase-like hydrolase (HAD) superfamily protein* (Jean-Baptiste et al., 2019; Wendrich et al., 2020; Graeff et al., 2021) and a *drought-responsive family protein (DIL9-3*) (Kim et al., 2021; Liu et al., 2022). CL5 (emb-3) was characterized by epidermal markers, including *3-ketoacyl-CoA synthase 20 (KCS20), which is* involved in cuticular wax biosynthesis (Suh et al., 2005; Wang et al., 2023), as well as *white-brown complex-like protein* linked to cutin metabolism (*ABCG*) (Panikashvili et al., 2010). CL6 (emb-4) was associated with leaf development, marked by a *YABBY family transcription factor*, which is also a known maize leaf marker (Wu et al., 2025). CL9 (emb-5) was linked to root, with markers such *as lateral root primordium* (*LRP*) (Singh et al., 2020), *aberrant root formation protein 4 (ALF4)* (DiDonato et al., 2004) and seed imbibition 2 (*SIP2*) (Jean-Baptiste et al., 2019).

In the scutellum-associated cluster (CL8, scu), key markers included *Late Embryogenesis Abundant protein (LEA-4-5), metallothionein-like protein*, and *glyoxalase I family protein (GLO1)*, all of which have been identified in the barley scutellum (Peirats-Llobet et al., 2023). CL7 (end-1) and CL11 (end-2) both represented residual endosperm. Marker genes supporting this included homologs of *kafirin PSKR2-like*, an endosperm storage protein (Li et al., 2024) and *alpha-zein 14*, a well-established maize endosperm marker (Wu et al., 2025). Two clusters were identified as proliferating meristematic cell populations, one corresponding to cells in the S phase and the other in the G2/M phase of the cell cycle. The G2/M phase cluster (CL12) was marked by *cyclin B2 (CCNB2), aurora1, and kinesin motor family* proteins, which have been identified as G2/M markers in the Arabidopsis shoot apex (Zhang et al., 2021). The S phase cluster (CL3, S) was characterized by the expression of *histones H1.3* and *H2AC12*, as well as *SMC3*, consistent with previously identified S-phase markers (Zhang et al., 2021). Clusters CL0 and CL2 contained the cells displaying a mixed and ambiguous transcriptional profile, and we were therefore unable to confidently assign it to any specific tissue or functional group at this stage.

In addition to described annotated groups, cluster 10 (B, 119 nuclei) did not fall into any of the above categories based on its transcriptional profile. Due to its distinct expression signature enriched for B chromosome-encoded genes, we refer to it as the cluster of nuclei comprising B chromosome. At this late developmental stage—approximately three weeks after fertilization—the B chromosome is retained in the limited number of cell. Given that only a fraction of nuclei in the embryo are derived from B chromosome containing proliferative tissue, the relative size of the B chromosome cluster (∼3% of all nuclei) is consistent with the expected proportion of those cells within the embryo.

### snRNA approach resolves the complexity of embryonic tissue with respect to the B chromosomes

In total, 35,745 genes were detected as expressed in the snRNA-seq dataset, including 3,024 genes located on the B chromosome. B chromosome-encoded genes were detected across all identified clusters; however, the number of B genes expressed per nucleus varied among clusters. Cells/nuclei in cluster 10 expressed higher number of genes encoded by the B chromosome (Figure EV2). This suggests that, while nuclei harbouring B chromosomes were distributed across all clusters, nearly all nuclei in cluster 10 carried the B chromosomes.

Cluster 10 was further characterized by the presence of 74 marker genes, of which 33 were located exclusively on the B chromosome (Source data 1). Of these 33 genes, 8 had been previously identified in an RNA-seq study as significantly upregulated in B chromosome possessing embryo (Bojdová et al., 2025).

This indicates that cluster 10 is distinctly enriched for B chromosome transcriptional signatures. Interestingly, the cluster 10 was not possible to align to any of the embryo tissue likely due to the B-cluster complexity. This assumption was confirmed *in silico*. Upon removing all B chromosome-encoded genes and subsequent clustering analysis, the B chromosome-associated cluster disappeared, and its constituent cells reassigned to embryonic and scutellar clusters (Figure EV1). This finding suggests that B chromosome expression overrides the typical tissue marker gene expression, indicating that the identity of this cluster is not tissue-specific but rather a consequence of high B chromosome transcriptional activity.

Single-nucleus RNA-seq outperformed whole embryo RNA-seq by detecting a larger number of expressed genes, including 3,024 located on the B chromosome, many of which were missed or underrepresented in conventional RNA-seq. Moreover, the transcriptional dominance of B chromosome marker genes caused B-positive nuclei to cluster together, independent of their tissue of origin. This clustering makes nuclei with B chromosomes easier to identify, enabling the targeted investigation of both known and novel B chromosome transcripts.

### Subclustering of B chromosome possessing cells reveals those undergoing elimination of B chromosome as well as novel candidate genes

While investigating the expression of previously identified 28 candidate genes associated with B chromosome elimination (Bojdová et al., 2025), we identified the B chromosome variant of *Structural Maintenance of Chromosomes 3* (*SMC3*) and Non-SMC *Condensin I Complex Subunit H* (*NCAPH*), as strong marker genes for the B cluster (CL10) (Source Data 1). SMC3, together with SMC1, forms the structural backbone of the cohesin ring complex, which encircles sister chromatids and prevents premature separation before anaphase. This cohesion is established during DNA replication in the S phase and persists until anaphase, when separase cleaves the cohesion complex to release the chromatids. NCAPH is a regulatory subunit of the condensin I complex, which is essential for chromosome condensation during mitosis and meiosis. Unlike the cohesin complex, which maintains chromatid cohesion, the condensin complex is responsible for compacting chromosomes to ensure their proper segregation. To better resolve transcriptional differences within this cluster, we performed independent clustering on nuclei contributing to CL10. This approach allowed us to focus on subtle variations within the B cluster (Fig. 3B).

**Figure 3:**
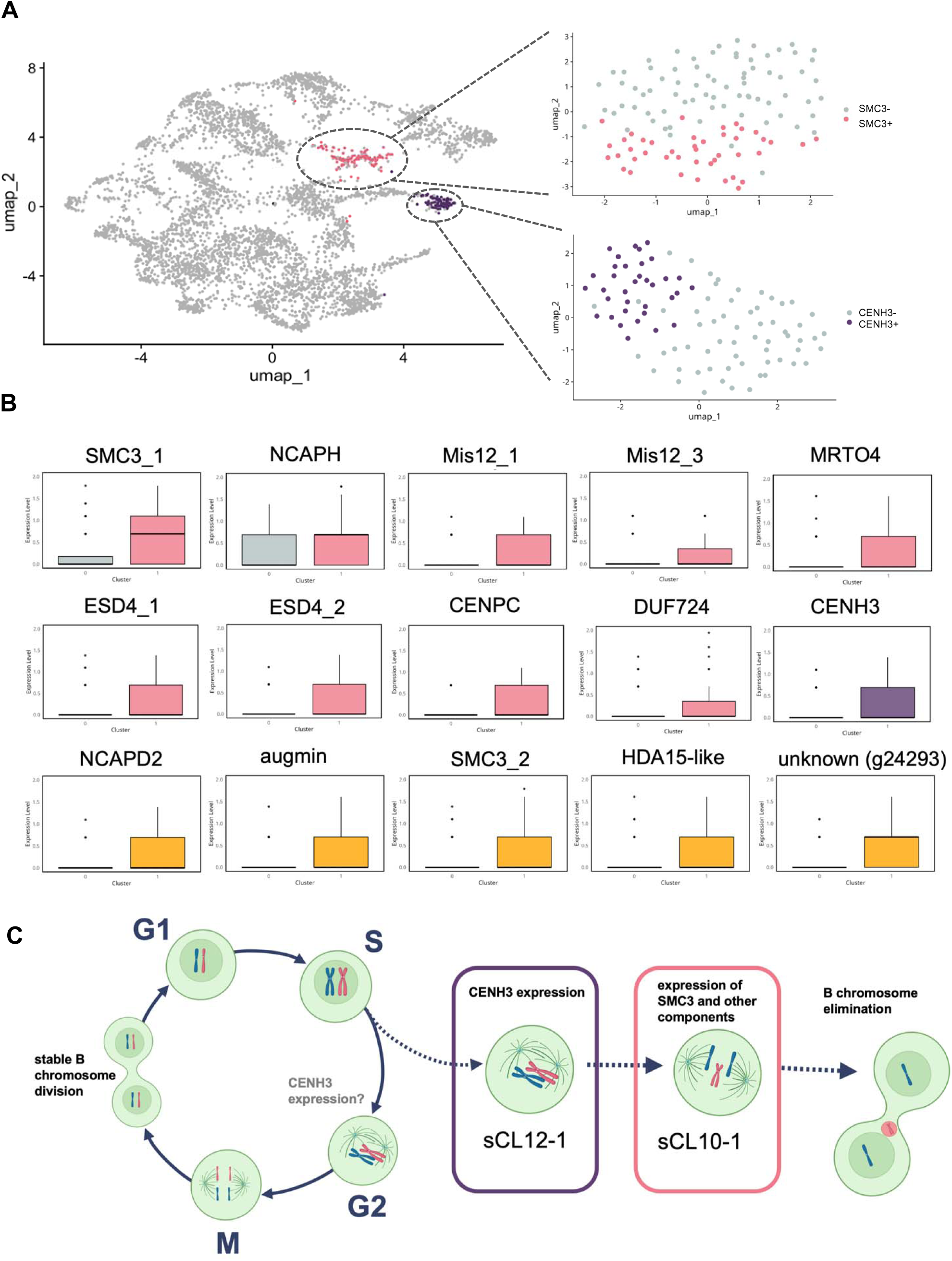
Analysis of two clusters associated with B chromosome elimination candidate gene expression. **A.** Left: UMAP representation of all captured single nuclei, as shown in Figure 2, highlighting the G2/M cluster (purple) and the B cluster (pink). Right: UMAP displaying independent clustering of these two clusters, with resolution set to 0.9. **B.** Boxplot depicting the expression levels of B chromosome elimination candidate genes in the B (purple, orange – novel candidates) and the G2/M (pink) cluster. Boxplots for candidate genes with no detectable expression were omitted. **C** Model of sequential gene expresion leading to the B chromosome elimination. The B chromosome shows either stable segregation during mitosis (left; no candidate is expressed) or is directed to elimination as indicated by changes in expression detected through analysis of sub-clusters of nuclei expressing B chromosome-encoded genes.

For the CL10-specific analysis, we applied the same Seurat processing pipeline as for the full dataset but adjusted the clustering resolution to 0.9. This resulted in the identification of a subcluster, sCL10-1, which was defined by 87 marker genes (Source data 3), including 38 genes localized to the B chromosome, among them *SMC3* and *NCAPH*. Additionally, sCL10-1 exhibited expression of other candidate genes encoded by the B chromosome—*CENP-C* and two copies of *Mis12* (*Mis12_1* and *Mis12_2*), which are involved in kinetochore function, an *mRNA turnover protein 4 homolog* (*MRTO4*), as well as *SUMO-specific protease ESD4*, and a *DUF724 domain-containing protein* (Fig. 3B). In contrast, subcluster sCL10-0 was defined by 63 marker genes, out of which only 1 was located on the B chromosome. Despite the lack of B chromosome-encoded genes among its top differentially expressed marker genes, sCL10-0 still showed clear expression of B chromosome genes. Considering the expression of B genes, both subclusters likely contain nuclei harbouring B chromosomes. However, the transcriptional profile of sCL10-0 lacks expression of candidate elimination-associated genes such as *Mis12* paralogs or *SMC3*, indicating that cells in this cluster may represent a population with stable B chromosomes. In contrast, sCL10-1 may correspond to a subset of B chromosome-containing cells actively undergoing, or primed for, the elimination process. Several additional marker genes of sCL10-1 followed the expression patterns of these B chromosome elimination candidates (Fig. 3B). Previously identified components of the condensin complex—subunits 2 and 3 (Bojdová et al., 2025)—were enriched of subunit 1 (*NCAPD2*; this study). More novel candidates, including *augmin subunit 3*, *histone deacetylase 15-like* (*HDA15-like*), and an additional copy of *SMC3*, showed the same expression pattern as well as numerous yet uncharacterized transcripts (Fig. 3B).

*SMC3*, *Mis12*, and condensin subunits have previously been associated with transcriptomic analyses of embryos undergoing B chromosome elimination in both *S. purpureosericeum* (Bojdová et al. 2025) and *Aegilops speltoides* (Boudichevskaia et al., 2020). Moreover, cohesion was previously discussed in the context of nondisjunction in rye (Banaei-Moghaddam et al., 2012; Chen et al., 2024), maize (Han et al., 2007) and *Aegilops speltoides* (Wu et al., 2019). Thus, identification of the same genes in our snRNA study suggests that these genes may contribute to elimination process.

Interestingly, sub-clustering within cluster CL10 clearly highlighted the advantage of the single-nucleus approach. In whole embryo RNA-seq analysis, which strictly distinguishes samples as +B and 0B, all differentially expressed genes are attributed to presence of the B chromosome. Contrary, the different transcriptional profiles of +B nuclei can be traced using snRNA-seq, indicating their different fates. While the nuclei in subcluster sCL10-0 may further maintain the B chromosome, the genes transcribed in cells contributing to sCL10-1 likely suggest on the onset of the elimination. This approach enabled us to track known elimination candidates and also to identify previously unrecognized ones within the eliminating subcluster sCL10-1.

### Transcription of B chromosome elimination-associated genes might be temporally separated

While investigating candidates for B chromosome elimination, we found that the B chromosome-specific variant of *CENH3* (*CENH3-B*), previously identified as a key candidate in transcriptomic analysis (Bojdová et al., 2025), was expressed in the G2/M cluster (CL12). Similarly to analysis of cluster CL10, we also applied an independent clustering approach to G2/M-phase cells at a resolution of 0.9 (Fig. 3A). This analysis identified a subcluster sCL12-1 defined by *CENH3-B* as one of marker genes. Notably, *CENH3-B* expression was absent in other cells within the G2/M cluster.

This observation raises the hypothesis that *CENH3-B* is expressed and potentially incorporated into nucleosomes only before/during specific processes, with B chromosome elimination being one such process. In contrast, histones of canonical nucleosomes are typically expressed and loaded during DNA replication in the S phase (Marzluff et al., 2008), which aligns with the observed expression of core histones in the S phase cluster (CL3). Meanwhile, CENH3 loading appears to be primarily restricted to G2 in both plants (Lermontova et al., 2006) and human (Badie et al., 2000). If CENH3-B functions as a determinant for B chromosome recognition during selective elimination, cells expressing CENH3-B and incorporating it into nucleosomes would be targeted for elimination in the subsequent division(s). Thus, the subcluster sCL12-1 likely represents a population of cells destined for future B chromosome elimination. Alternatively, B chromosome-encoded CENH3-B might be incorporated in each G2 phase to maintain the B chromosome identity.

Considering that both genes *SMC3-B* and *CENH3-B* may contribute to the B chromosome elimination, the splitting of the nuclei expressing these genes is surprising, alike the absence of the nuclei co-expressing both genes. In our data, we detected only two nuclei positive for the transcripts of both genes. Fact that majority of nuclei expressed exclusively one of these two key candidate genes implies that the transcription of the individual genes driving B chromosome to elimination is likely step-wise. CENH3-B might be first incorporated into centromeric nucleosomes during G2, marking B chromosomes for recognition. Subsequently, during mitosis, the expression of SMC3-B and other components may facilitate aberrant cohesion or segregation, ultimately leading to B chromosome elimination. This sequential model, illustrated in Figure 3C, highlights the temporal coordination of CENH3-B loading and SMC3-B expression as potential determinants of B chromosome elimination.

Based on snRNA-seq data analysis in *Sorghum purpureosericeum* embryos, this study for the first time provided insights into the process of B chromosome elimination at the single-nucleus level. This approach captured substantially more B chromosome transcripts than conventional RNA-seq and revealed that their expression is so dominant that nuclei harbouring B chromosomes cluster together. Further, it suggested on the identity of the cells in which the B chromosome is eliminated and identified additional candidate genes potentially involved in the process. Finally, it uncovered the sequential timing of the expression of key genes that may be involved the B chromosome elimination. This study thus demonstrates the utility of the snRNA-seq method for unravelling the complexity of embryonic tissue with respect to the B chromosomes elimination and lays the groundwork for future single-cell investigations in B chromosome biology.

## Methods

### Plant material and cultivation

The progeny of *Sorghum purpureosericeum* plants possessing B chromosomes (+B) harvested at Institute of Experimental Botany (Czech Republic) was used as plant material. Plants were grown under short-day conditions, featuring 10 hours of daylight, with daytime temperatures maintained at 29 °C and nighttime temperatures at 25 °C. The presence of the B chromosome in plants was scored in initial spike using flow cytometry, following the methodology published previously (Karafiátová et al., 2021).

### Embryo harvesting, sectioning and in situ hybridization

All embryos were collected from +B plants. The stage of embryo development was determined as described in Bojdová et al., (2025). Embryos at stage of ∼ 21 DAP were further screened for B chromosome presence, which was assessed in endosperm using B chromosome-specific PCR marker SpuB (Karafiátová et al., 2024). +B embryos were selected for sectioning and snRNA-seq.

Simultaneously, two protocols for embryo-sectioning were employed. First, wax-sectioning coupled with toluidine blue staining was used to visualize anatomy of the embryos. Second, the localization of the B chromosome in embryonic tissues was detected by fluorescent *in situ* hybridization on embryo cryo-sections. Both protocols for sectioning together with probe labelling and FISH followed the methodologies described in Karafiátová et al. (2024) and Bojdová et al. (2025).

### Preparation of nuclei suspension and flow cytometric purification

About 60 deep frozen +B embryos were pooled and used for nuclei isolation. Sorghum nuclei suspension was prepared using Singulator^TM^ 100 instrument (S2 Genomics, Livermore, CA, USA) according to pre-designed protocol for low volume nuclei with the optimized settings. Briefly, the frozen embryos were transferred into the Singulator cartridge and soaked in 1.5mL NIS reagent (nuclei isolation solution, delivered with Singulator machine) supplemented with 1,500 U of RNase inhibitor (cat.no. 03335402001, Roche, Basel, Swiss). Embryos homogenization was performed in Singulator instrument cooled down to 4°C applying single-shot mode with the setting as follows: auto-mince ON, incubation time - 10 min, mixing type-top, disruption type – default, incubation temperature – cold, mixing speed – fast, disruption speed – fast. The clean nuclei suspension was centrifuged for 5 min at 500g and 4°C, supernatant discarded and pellet resuspended in 1mL of NIS containing 1,000 U of RNase inhibitor. Finally, nuclei were filtered through 20µm mesh, DAPI stained (2mg/mL) and analyzed using flow cytometer FACSAria SORP (BD Biosciences, San Jose, CA, USA). In total, 100,000 of G1, S and G2 population of embryonic nuclei (containing both +B and 0B nuclei) were purified into 20µl of 1xPBS with BSA (40µg/mL) and 50 U of RNase inhibitor resulting in the suspension concentration of approx. 830 nuclei/µl.

### Single-nuclei RNA-seq library preparation and sequencing

Sequencing library was prepared using Chromium Next GEM Single Cell 3’v3.1. (10x Genomics, Pleasanton, CA, USA) according to manufactureŕs instructions with minor modification. Shortly, a master mix containing of about 17,000 nuclei were loaded into the Chromium chip and partitioning of individual nuclei was performed in Chromium Controller (10x Genomics). Before sequencing library construction, cDNA was amplified in 12 PCR cycles, yielding ∼ 40 ng of amplified cDNA. After fragmentation and adaptor ligation, final libraries were prepared using the Dual Index Kit TT, Set A (10x Genomics) in 15 PCR cycles. The quality of the final library was controlled by capillary electrophoresis on Bionalyzer 2100 (Agilent) using Agilent High Sensitivity DNA chip. The library was sequenced on Illumina NovaSeq 6000.

### snRNA-seq data mapping and post-processing

snRNA-seq data were analyzed using the *S. purpureosericeum* reference sequence and annotation published recently (Bojdová et al., 2025). The annotation was further modified with peaks2UTR pipeline, which corrected missing 3’UTR to increase number of detected genes (Haese-Hill et al., 2023). Reads were mapped to the reference sequence using STAR 2.7.11b. The maximum intron length was set to 50,000 bases and multimapping limited to 15 loci. Cell barcodes and UMIs were processed using soloType CB_UMI_Simple with a 16 bp cell barcode (--soloCBstart 1 --soloCBlen 16) and a 12 bp UMI (--soloUMIstart 17 --soloUMIlen 12). UMI deduplication was performed using 1MM_Directional, and multi-gene UMIs were filtered (MultiGeneUMI). Reads were output as a sorted BAM file, and count matrices (features, barcodes, matrix) were generated using --soloFeatures GeneFull. We applied emptyDrops74 from the DropletUtils package (version 1.6.1) to distinguish true cells from empty droplets (Griffiths et al., 2018; Lun et al., 2019).

For subsequent analysis, Seurat package (v.5.1.0) was used (Hao et al., 2024; Hao et al., 2021; Stuart et al., 2019). Data were normalized using the SCTransform function (parameters: return.only.var.genes = TRUE, method = "glmGamPoi“, variable.features.n = NULL) (Hafemeister & Satija, 2019). Dimensionality reduction was performed with RunPCA on all genes, and the top 30 principal components were used for Uniform Manifold Approximation and Projection (UMAP) and clustering of nuclei.

### Clustering and cell-type annotation

Generation of spot clusters was made using the ‘FindNeighbors’ (settings: reduction = ‘pca’, dims = 1:30) followed by the ‘FindClusters’ functions in Seurat. Cluster resolution was set to 0.5. To identify marker genes for each cluster, pair-wise comparisons of individual clusters against all other clusters were performed using the ‘FindAllMarkers’ function (settings: min.pct = 0.03, only.pos = T). To characterize marker genes, we searched for orthologous genes in maize, barley and sorghum at https://plants.ensembl.org/ and annotated embryo cell types using previously curated marker transcripts resulting from scRNA-seq analyses (He et al., 2024). To inspect clusters B and G2/M in detail, we subsetted the B or G2/M cells from SCT normalized Seurat object and performed the pipeline from RunPCA as previously described. The resolution was set to 0.9 based upon closer examination of cluster trees.

## Acknowledgements

Computational resources were provided by the e-INFRA CZ project (ID:90254), supported by the Ministry of Education, Youth and Sports of the Czech Republic and by the ELIXIR-CZ project (ID:90255), part of the international ELIXIR infrastructure. We acknowledge the support of the GeneCore Facility at the Institute of Biotechnology of the Czech Academy of Sciences for help with snRNA-Seq library preparation. Authors thank to S2 Genomics (Livermore, CA, USA) for lending them Singulator^TM^ 100 instrument for nuclei preparation.

## Data availability

Raw snRNA-seq reads are available from the European Nucleotide Archive (ENA) under accession PRJEB97401. Genome assembly originally published in Bojdová et al. (2025) is available in ENA under ID - GCA_964647645.

## References

Badie, C., Itzhaki, J. E., Sullivan, M. J., Carpenter, A. J., & Porter, A. C. G. (2000). Repression of CDK1 and Other Genes with CDE and CHR Promoter Elements during DNA Damage-Induced G 2 /M Arrest in Human Cells. Molecular and Cellular Biology, 20(7), 2358–2366. 10.1128/MCB.20.7.2358-2366.2000

Baenziger, H. (1962). Supernumerary chromosomes in diploid and tetraploid forms of crested wheatgrass. Canadian Journal of Botany, 40(4), 549–561. 10.1139/b62-052

Banaei-Moghaddam, A. M., Schubert, V., Kumke, K., Weiβ, O., Klemme, S., Nagaki, K., Macas, J., González-Sánchez, M., Heredia, V., Gómez-Revilla, D., González-García, M., Vega, J. M., Puertas, M. J., & Houben, A. (2012). Nondisjunction in Favor of a Chromosome: The Mechanism of Rye B Chromosome Drive during Pollen Mitosis. The Plant Cell, 24(10), 4124–4134. 10.1105/tpc.112.105270

Becht, E., McInnes, L., Healy, J., Dutertre, C. -A., Kwok, I. W. H., Ng, L. G., Ginhoux, F., & Newell, E. W. (2019). Dimensionality reduction for visualizing single-cell data using UMAP. Nature Biotechnology, 37(1), 38–44. 10.1038/nbt.4314

Bojdová, T., Hloušková, L., Holušová, K., Svačina, R., Hřibová, E., Ilíková, I., Thiel, J., Kim, G., Pleskot, R., Houben, A., Bartoš, J. and Karafiátová, M. (2025). Sorghum embryos undergoing B chromosome elimination express B-variants of mitotic-related genes. bioRxiv doi:10.1101/2025.09.30.679447

Borodin, P., Chen, A., Forstmeier, W., Fouché, S., Malinovskaya, L., Pei, Y., Reifová, R., Ruiz-Ruano, F. J., Schlebusch, S. A., Sotelo-Muñoz, M., Torgasheva, A., Vontzou, N., & Suh, A. (2022). Mendelian nightmares: the germline-restricted chromosome of songbirds. Chromosome Research, 30(2-3), 255–272. 10.1007/s10577-022-09688-3

Boudichevskaia, A., Ruban, A., Thiel, J., Fiebig, A., & Houben, A. (2020). Tissue-Specific Transcriptome Analysis Reveals Candidate Transcripts Associated with the Process of Programmed B Chromosome Elimination in Aegilops speltoides. International Journal of Molecular Sciences, 21(20). 10.3390/ijms21207596

Chen, J., Bartoš, J., Boudichevskaia, A., Voigt, A., Rabanus-Wallace, M. T., Dreissig, S., Tulpová, Z., Šimková, H., Macas, J., Kim, G., Buhl, J., Bürstenbinder, K., Blattner, F. R., Fuchs, J., Schmutzer, T., Himmelbach, A., Schubert, V., & Houben, A. (2024). The genetic mechanism of B chromosome drive in rye illuminated by chromosome-scale assembly. Nature Communications, 15(1). 10.1038/s41467-024-53799-w

D’Ambrosio, U., Pilar Alonso-Lifante, M., Barros, K., Kovařík, A., Mas de Xaxars, G., & Sònia Garcia, G. (2017). B-chrom: a database on B-chromosomes of plants, animals and fungi. New Phytologist, 216, 635–642. 10.1111/nph.14723

DiDonato, R. J., Arbuckle, E., Buker, S., Sheets, J., Tobar, J., Totong, R., Grisafi, P., Fink, G. R., & Celenza, J. L. (2004). Arabidopsis ALF4 encodes a nuclear-localized protein required for lateral root formation. The Plant Journal, 37(3), 340–353. 10.1046/j.1365-313X.2003.01964.x

Graeff, M., Rana, S., Wendrich, J. R., Dorier, J., Eekhout, T., Aliaga Fandino, A. C., Guex, N., Bassel, G. W., De Rybel, B., & Hardtke, C. S. (2021). A single-cell morpho-transcriptomic map of brassinosteroid action in the Arabidopsis root. Molecular Plant, 14(12), 1985–1999. 10.1016/j.molp.2021.07.021

Griffiths, J. A., Richard, A. C., Bach, K., Lun, A. T. L., & Marioni, J. C. (2018). Detection and removal of barcode swapping in single-cell RNA-seq data. Nature Communications, 9(1). 10.1038/s41467-018-05083-x

Haese-Hill, W., Crouch, K., Otto, T. D., & Marschall, T. (2023). Peaks2utr: a robust Python tool for the annotation of 3′ UTRs. Bioinformatics, 39(3). 10.1093/bioinformatics/btad112

Hafemeister, C., & Satija, R. (2019). Normalization and variance stabilization of single-cell RNA-seq data using regularized negative binomial regression. Genome Biology, 20(1). 10.1186/s13059-019-1874-1

Han, F., Gao, Z., Yu, W., & Birchler, J. A. (2007). Minichromosome Analysis of Chromosome Pairing, Disjunction, and Sister Chromatid Cohesion in Maize. The Plant Cell, 19(12), 3853–3863. 10.1105/tpc.107.055905

Hao, Y., Hao, S., Andersen-Nissen, E., Mauck, W. M., Zheng, S., Butler, A., Lee, M. J., Wilk, A. J., Darby, C., Zager, M., Hoffman, P., Stoeckius, M., Papalexi, E., Mimitou, E. P., & Jain, J. (2021). Integrated analysis of multimodal single-cell data. Cell, 184(13), 3573–3587.e29. 10.1016/j.cell.2021.04.048

Hao, Y., Stuart, T., Kowalski, M. H., Choudhary, S., Hoffman, P., Hartman, A., Srivastava, A., Molla, G., Madad, S., Fernandez-Granda, C., & Satija, R. (2024). Dictionary learning for integrative, multimodal and scalable single-cell analysis. Nature Biotechnology, 42(2), 293–304. 10.1038/s41587-023-01767-y

He, Z., Luo, Y., Zhou, X., Zhu, T., Lan, Y., & Chen, D. (2024). ScPlantDB: a comprehensive database for exploring cell types and markers of plant cell atlases. Nucleic Acids Research, 52(D1), D1629–D1638. 10.1093/nar/gkad706

Hodson, C. N., Ross, L., & Dion-Côté, A. -M. (2021). Evolutionary Perspectives on Germline-Restricted Chromosomes in Flies (Diptera). Genome Biology and Evolution, 13(6). 10.1093/gbe/evab072

Jean-Baptiste, K., McFaline-Figueroa, J. L., Alexandre, C. M., Dorrity, M. W., Saunders, L., Bubb, K. L., Trapnell, C., Fields, S., Queitsch, C., & Cuperus, J. T. (2019). Dynamics of Gene Expression in Single Root Cells of Arabidopsis thaliana. The Plant Cell, 31(5), 993–1011. 10.1105/tpc.18.00785

Kao, P., Schon, M. A., Mosiolek, M., Enugutti, B., & Nodine, M. D. (2021). Gene expression variation in Arabidopsis embryos at single-nucleus resolution. Development, 148(13). 10.1242/dev.199589

Karafiátová, M., Bednářová, M., Said, M., Čížková, J., Holušová, K., Blavet, N., & Bartoš, J. (2021). The B chromosome of Sorghum purpureosericeum reveals the first pieces of its sequence. Journal of Experimental Botany, 72(5), 1606–1616. 10.1093/jxb/eraa548

Karafiátová, M., Bojdová, T., Stejskalová, M., Harnádková, N., Kumar, V., Houben, A., Chen, J., Doležalová, A., Honys, D., & Bartoš, J. (2024). Unravelling the unusual: chromosome elimination, nondisjunction and extra pollen mitosis characterize the B chromosome in wild sorghum. New Phytologist, 243(5), 1840–1854. 10.1111/nph.19954

Kato, T., Morita, M. T., Fukaki, H., Yamauchi, Y., Uehara, M., Niihama, M., & Tasaka, M. (2002). SGR2, a Phospholipase-Like Protein, and ZIG/SGR4, a SNARE, Are Involved in the Shoot Gravitropism of Arabidopsis. The Plant Cell, 14(1), 33–46. 10.1105/tpc.010215

Kim, J. -Y., Symeonidi, E., Pang, T. Y., Denyer, T., Weidauer, D., Bezrutczyk, M., Miras, M., Zöllner, N., Hartwig, T., Wudick, M. M., Lercher, M., Chen, L. -Q., Timmermans, M. C. P., & Frommer, W. B. (2021). Distinct identities of leaf phloem cells revealed by single cell transcriptomics. The Plant Cell, 33(3), 511–530. 10.1093/plcell/koaa060

Lermontova, I., Schubert, V., Fuchs, J., Klatte, S., Macas, J., & Schubert, I. (2006). Loading of Arabidopsis Centromeric Histone CENH3 Occurs Mainly during G2 and Requires the Presence of the Histone Fold Domain. The Plant Cell, 18(10), 2443–2451. 10.1105/tpc.106.043174

Li, X., Liu, W., Wang, G., Sun, S. S. -M., Yuan, L., & Wang, J. (2024). Improving digestibility of sorghum proteins by CRISPR /Cas9-based genome editing. Food and Energy Security, 13(1). 10.1002/fes3.506

Liu, W., Zhang, Y., Fang, X., Tran, S., Zhai, N., Yang, Z., Guo, F., Chen, L., Yu, J., Ison, M. S., Zhang, T., Sun, L., Bian, H., Zhang, Y., & Yang, L. (2022). Transcriptional landscapes of de novo root regeneration from detached Arabidopsis leaves revealed by time-lapse and single-cell RNA sequencing analyses. Plant Communications, 3(4). 10.1016/j.xplc.2022.100306

Long, Y., Liu, Z., Jia, J., Mo, W., Fang, L., Lu, D., Liu, B., Zhang, H., Chen, W., & Zhai, J. (2021). FlsnRNA-seq: protoplasting-free full-length single-nucleus RNA profiling in plants. Genome Biology, 22(1). 10.1186/s13059-021-02288-0

Lun, A. T. L., Riesenfeld, S., Andrews, T., Dao, T. P., Gomes, T., & Marioni, J. C. (2019). EmptyDrops: distinguishing cells from empty droplets in droplet-based single-cell RNA sequencing data. Genome Biology, 20(1). 10.1186/s13059-019-1662-y

Marzluff, W. F., Wagner, E. J., & Duronio, R. J. (2008). Metabolism and regulation of canonical histone mRNAs: life without a poly(A) tail. Nature Reviews Genetics, 9(11), 843–854. 10.1038/nrg2438

Mendelson, D., & Zohary, D. (1972). Behaviour and transmission of supernumerary chromosomes in Aegilops speltoides. Heredity, 29(3), 329–339. 10.1038/hdy.1972.97

Mochizuki, A. (1957). B chromosomes in Aegilops mutica Boiss. Wheat Information Service, 5, 9–11.

Nygren, A. (1957). Poa timoleontis Heldr., a new diploid species of the section Bolbophorum and Gr. with accessory chromosomes only in meiosis. LantbrHögsk. Annual Report Agricultural College of Sweden, 23, 489–495.

Panikashvili, D., Shi, J. X., Bocobza, S., Franke, R. B., Schreiber, L., & Aharoni, A. (2010). The Arabidopsis DSO/ABCG11 Transporter Affects Cutin Metabolism in Reproductive Organs and Suberin in Roots. Molecular Plant, 3(3), 563–575. 10.1093/mp/ssp103

Peirats-Llobet, M., Yi, C., Liew, L. C., Berkowitz, O., Narsai, R., Lewsey, M. G., & Whelan, J. (2023). Spatially resolved transcriptomic analysis of the germinating barley grain. Nucleic Acids Research, 51(15), 7798–7819. 10.1093/nar/gkad521

Ruban, A., Schmutzer, T., Wu, D. D., Fuchs, J., Boudichevskaia, A., Rubtsova, M., Pistrick, K., Melzer, M., Himmelbach, A., Schubert, V., Scholz, U., & Houben, A. (2020). Supernumerary B chromosomes of Aegilops speltoides undergo precise elimination in roots early in embryo development. Nature Communications, 11(1). 10.1038/s41467-020-16594-x

Singh, S., Yadav, S., Singh, A., Mahima, M., Singh, A., Gautam, V., & Sarkar, A. K. (2020). Auxin signaling modulates LATERAL ROOT PRIMORDIUM 1 (LRP 1) expression during lateral root development in Arabidopsis. The Plant Journal, 101(1), 87–100. 10.1111/tpj.14520

Stuart, T., Butler, A., Hoffman, P., Hafemeister, C., Papalexi, E., Mauck, W. M., Hao, Y., Stoeckius, M., Smibert, P., & Satija, R. (2019). Comprehensive Integration of Single-Cell Data. Cell, 177(7), 1888–1902.e21. 10.1016/j.cell.2019.05.031

Suh, M. C., Samuels, A. L., Jetter, R., Kunst, L., Pollard, M., Ohlrogge, J., & Beisson, F. (2005). Cuticular Lipid Composition, Surface Structure, and Gene Expression in Arabidopsis Stem Epidermis. Plant Physiology, 139(4), 1649–1665. 10.1104/pp.105.070805

Wang, Y., Liu, Y., Pan, X., Wan, Y., Li, Z., Xie, Z., Hu, T., & Yang, P. (2023). A 3-Ketoacyl-CoA Synthase 10 (KCS10) Homologue from Alfalfa Enhances Drought Tolerance by Regulating Cuticular Wax Biosynthesis. Journal of Agricultural and Food Chemistry, 71(40), 14493–14504. 10.1021/acs.jafc.3c03881

Wendrich, J. R., Yang, B. J., Vandamme, N., Verstaen, K., Smet, W., Van de Velde, C., Minne, M., Wybouw, B., Mor, E., Arents, H. E., Nolf, J., Van Duyse, J., Van Isterdael, G., Maere, S., & Saeys, Y. (2020). Vascular transcription factors guide plant epidermal responses to limiting phosphate conditions. Science, 370(6518). 10.1126/science.aay4970

Wu, D. D., Ruban, A., Fuchs, J., Macas, J., Novák, P., Vaio, M., Zhou, Y. H., & Houben, A. (2019). Nondisjunction and unequal spindle organization accompany the drive of Aegilops speltoides B chromosomes. New Phytologist, 223(3), 1340–1352. 10.1111/nph.15875

Wu, H., Zhang, R., Niklas, K. J., & Scanlon, M. J. (2025). Multiplexed transcriptomic analyzes of the plant embryonic hourglass. Nature Communications, 16(1). 10.1038/s41467-024-55803-9

Zhang, T. -Q., Chen, Y., & Wang, J. -W. (2021). A single-cell analysis of the Arabidopsis vegetative shoot apex. Developmental Cell, 56(7), 1056–1074.e8. 10.1016/j.devcel.2021.02.021

Zhao, Y., Hu, Y., Dai, M., Huang, L., & Zhou, D. -X. (2009). The WUSCHEL-Related Homeobox Gene WOX11 Is Required to Activate Shoot-Borne Crown Root Development in Rice. The Plant Cell, 21(3), 736–748. 10.1105/tpc.108.061655

