## Supplementary Figures for "Single-nucleus transcriptome analysis provides new insights into B chromosome elimination in sorghum"

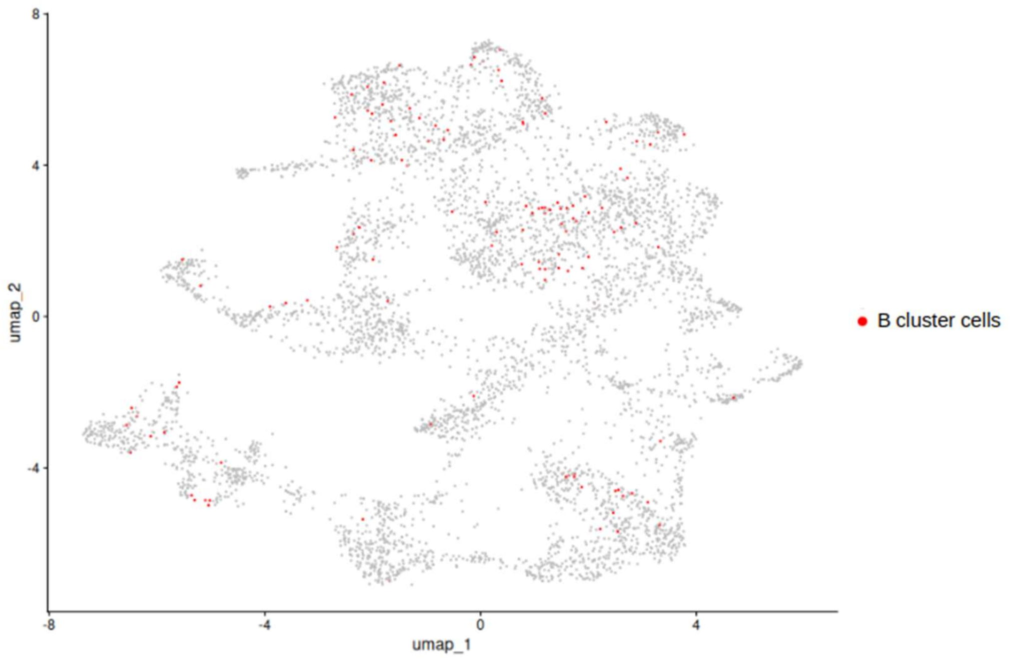


**Figure EV1:** Reclustering of snRNA-seq data after removal of B chromosome-encoded genes. Cells from the original B chromosome-associated cluster (highlighted in red) are redistributed into embryonic and scutellar clusters, confirming that this cluster was driven by B chromosome transcription rather than tissue identity.

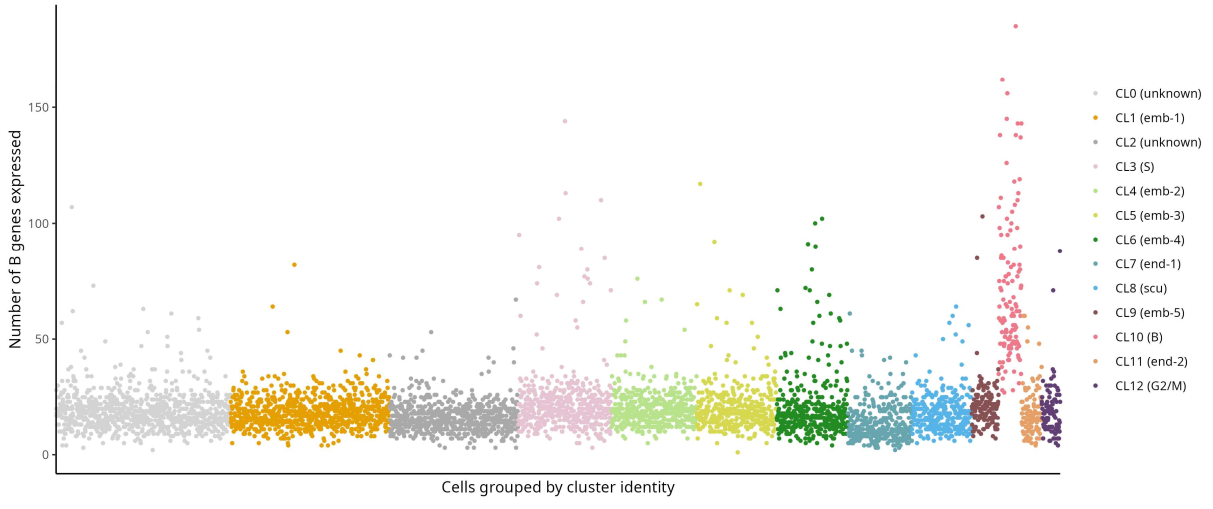


**Figure EV2:** Expression of high confidence B chromosome genes across single nuclei grouped by cluster identity. Cells are ordered by their assigned clusters, and the number of B chromosome-encoded genes expressed is shown per nucleus. While most clusters display low to moderate B gene expression, cluster 10 (B) is markedly enriched, with many nuclei expressing a high number of B genes.
